# Forward genetic screens identified mutants with defects in trap morphogenesis in the nematode-trapping fungus *Arthrobotrys oligospora*

**DOI:** 10.1101/2020.09.15.298422

**Authors:** Tsung-Yu Huang, Yi-Yun Lee, Guillermo Vidal-Diez de Ulzurrun, Yen-Ping Hsueh

**Author notes:** Corresponding author: Institute of Molecular Biology, Academia Sinica, 128 Academia Road, Section 2, Nangang, Taipei, 115 Taiwan.

## Abstract

Nematode-trapping fungi (NTF) are carnivorous fungi that prey on nematodes under nutrient-poor conditions via specialized hyphae that function as traps. The molecular mechanisms involved in the interactions between nematode-trapping fungi and their nematode prey are largely unknown. In this study, we conducted forward genetic screens to identify potential genes and pathways that are involved in trap morphogenesis and predation in the NTF *Arthrobotrys oligospora.* Using Ethyl methanesulfonate and UV as the mutagens, we generated 5552 randomly-mutagenized *A. oligospora* strains and identified 15 mutants with strong defects in trap morphogenesis. Whole genome sequencing and bioinformatic analyses revealed mutations in genes with roles in signaling, transcription or membrane transport that may contribute to the defects of trap morphogenesis in these mutants. We further conducted functional analyses on a candidate gene, *YBP-1*, and demonstrate that mutation of that gene was causative of the phenotypes observed in one of the mutants. The methods established in this study might provide helpful insights for establishing forward genetic screening methods for other non-model fungal species.

## Introduction

Model organisms have contributed enormously to unraveling the fundamental principles of biology. However, in some cases, model organisms might not be the most ideal investigatory system and studies on non-model species could better answer specific questions (Russell et al., 2017). Fortunately, by adopting increasingly more affordable and advanced next-generation sequencing technologies, intensive study at molecular and cellular levels of non-model species is now possible (Ellegren and evolution, 2014).

The fungal kingdom harbors rich biodiversity, with an estimated ~12 million species (Wu et al., 2019). Apart from for several model yeast and filamentous species, particularly pathogenic ones, we have very limited molecular or cellular knowledge on the vast majority of fungi. Fungi occur in essentially all habitats on Earth, and many species have evolved unique traits. For example, many species of Orbiliaceae (Ascomycota) are predatory fungi that prey on nematodes when local nutrients are scarce by means of specialized mycelial structures. More than 200 of these nematode-trapping fungi (NTF) have been described to date, with *Arthrobotrys oligospora* being the best studied species (Nordbring-Hertz et al., 2011). *A. oligospora* is known to eavesdrop on nematode ascaroside pheromones to trigger trap morphogenesis and to produce volatile compounds mimicking food and sex cues that lure nematodes, and predatory abilities among natural populations of this species can vary considerably (Hsueh et al., 2017; Hsueh et al., 2013; Yang et al., 2020).

NTF hold great promise as biocontrol agents to combat plant-parasitic nematodes, which have been estimated to cause USD$ 80 million in crop loss annually (Gray, 1983; Lopez-Llorca et al., 2007; Wang et al., 2015). However, since we know little about their biology and the cellular/molecular mechanisms governing the switch from saprophytic to predatory lifestyles, the full nematicidal potential of NTF cannot yet be harnessed. Nematode-derived cues are key signals triggering the switch to a predatory lifestyle switch in NTF (Hsueh et al., 2013). Consequently, signaling pathways could be anticipated to play vital roles in this lifestyle switch. Indeed, a few such genes have been demonstrated as necessary for trap morphogenesis in *A. oligopspora*. These include the mitogen-activated protein kinase (MAPK), Slt2, which is involved in the cell wall integrity signaling pathway in yeast (Zhen et al., 2018), a pH-sensing receptor, palH, that functions in the pH signal transduction pathway, and a NADPH oxidase, NoxA, that controls ROS signaling responses in *A. oligospora* (Li et al., 2019; Li et al., 2017). More recently, it has been demonstrated that the G protein beta subunit, Gpb1, is required for the predatory lifestyle transition in *A. oligospora* (Yang et al., 2020). Nevertheless, more intensive study is needed to gain further insight into the mechanisms governing these inter-kingdom predator-prey interactions.

In this study, we have established methods for conducting forward genetic screens of *A. oligospora*. By applying this approach together with whole genome sequencing, we demonstrated that it is possible to identify genes playing important roles in trap morphogenesis and, further, to unveil the causative mutation in a mutant that failed to develop traps without the need to conduct genetic crosses. Our work illustrates that it is feasible to conduct random mutagenesis on a fungal species for which the sexual cycle has not been well studied in the laboratory and to identify causative mutations for phenotypic traits. Through our forward genetic screens, we have identified novel players necessary for the predatory lifestyle of *A. oligospora*.

## Materials and methods

### Strains, media, culture conditions

*Arthrobotrys oligospora* strain TWF154 was used in this study, a wild isolate we have sampled in Taiwan. Genomic data on this strain can be accessed from the National Center for Biotechnology Information GenBank under accession number SOZJ00000000 (Yang et al., 2020). All corresponding knockout mutants were obtained in a *ku70* strain background. Our use of *ku70* protoplasts increased the efficiency of targeted gene knockout as cells may fail to enter the nonhomologous end-joining pathway without Ku70 protein, resulting in greater likelihood of undergoing homologous recombination during DNA repair.

We used potato dextrose agar (PDA) and low nutrient medium (LNM) as fungal solid media, whereas yeast nitrogen base without amino acids (YNB) and potato dextrose broth (PDB) acted as liquid media. We used *Caenorhabditis elegans* wild-type strain N2 as nematode prey, which were maintained on nematode growth media (NGM) plates with *Escherichia coli* OP50 as food. All cultures were incubated at 25 °C.

### Mutagenesis

TWF154 was cultured on PDA for 5 days and the hyphae and spores were collected into 50 ml PDB for liquid culture. After culturing for 2 days at 25 °C, the liquid culture was blended and then treated with 10 ml Vino Taste Pro (80 mg/ml in MN buffer) and chitinase for 8-10 h at 30 °C and 200 rpm in an incubator to digest fungal cell walls. Digested cells were then filtered through two layers of sterile miracloth (EMD Millipore) and washed with sterile STC buffer (1.2 M D-Sorbitol, 10 mM Tris-HCl (pH 7.5), 50 mM CaCl_2_).

Next, we spread 5×10^4^ protoplasts onto a regeneration plate and subjected them to a treatment of either 6 sec of 15 W UV or 12 μg/ml ethyl methanesulfonate (EMS) to cause random mutations in the genomes. Plates were then cultured under dark conditions at 25 °C to prevent DNA repair. Any colonies formed were then further inoculated onto PDA 48-well plates.

### Genetic screen on mutants with trapping defects

To screen out mutants exhibiting defects in trapping *C. elegans*, colonies that grew after mutagenesis were inoculated onto LNM 48-well plates. Colonies in each well were then exposed overnight to thirty specimens of N2, and those exhibiting weak trapping ability after 24 hours were selected for rescreening. The rescreening process was conducted on 5-cm LNM plates on which mutant fungal lines were exposed to ~100 N2. By rescreening we could exclude false positive mutants, resulting in more accurate phenotyping of trapping defects.

### Trap quantification

To quantify trap formation, fungal strains were inoculated onto fresh 3-cm LNM plates and grown for 3 days. Then, thirty L4 larval-stage N2 were added to the plate and washed away after 6 hours. Fungal cultures were incubated at 25 °C and we calculated number of traps from half of the plate after 24 hours and presented the results in traps/cm^2^.

### Visualization of trap morphology

Fungal strains were inoculated onto 12-well LNM plates, and 0.1% SCRI Renaissance 2200 (SR2200; a dye that binds to beta-1,3-glucan) was added to the medium. Thirty L4 larval-stage N2 were added to the plates and 24-h later the plates were imaged at 40x magnification using an Axio Observer Z1 system.

### Whole-genome sequencing analysis

Genomic DNA extracted from 16 mutants was subjected to whole genome sequencing using an Illumina sequencing system. Approximately 18 million reads were trimmed to generate a 250 base-pair (bp) library and a paired-end sequencing protocol enabled us to derive more accurate sequencing results.

For data analysis, index files from the TWF154 reference genome (Yang et al., 2020) were created using samtools (Li, 2011) and bwa (Li and Durbin, 2009). Then, we trimmed the adaptors with Trimmomatic (Bolger et al., 2014) and filtered out low-quality reads from each of the sequenced mutants. Trimmed reads from each mutant were aligned to the reference genome using bwa-mem (Li, 2013) and converted to BAM files. Next, we used Picard (Picard toolkit., 2019) to identify duplicates, and GATK (Poplin et al., 2017) was employed for SNV (single nucleotide variation) and INDEL (insertion/deletion) calling in each file. Two separate files of all variants were then generated, one of which contained only INDELs and the other only SNPs. To focus on the most relevant mutations, the SNP and INDEL files were filtered using gatk VariantFiltration and gatk SelectVariants (Poplin et al., 2017) with the following criteria: QD <2.0, MQ < 40.0, QUAL < 100, MQRankSum < −12.5, SOR > 4.0, FS > 60.0, ReadPosRankSum < −8.0. The mutations were annotated in ANNOVAR (Wang et al., 2010) by comparing the files to the reference genome. To narrow down potential candidate genes, we focused on exonic regions and excluded synonymous mutations as well as mutations occurring more than twice among the mutants. Genes potentially contributing to trapping defects were validated by gene ontology prediction.

### Transformation

We placed 10^6^ protoplasts on ice in a 50-ml centrifuge tube for 30 min with 5 μg of knockout cassette DNA (see below). Then, five volumes of PTC buffer (40% polyethylene glycol 3350, 10 mM Tris-HCl (pH 7.5), 50 mM CaCl_2_) was added and gently mixed by inverting the tube. After multiple inversions, the tube was kept at room temperature for 20 min. Lastly, protoplasts were mixed with regeneration agar (3% acid-hydrolize casein, 3% yeast extract, 0.5 M sucrose, 10% agar) and 200 μg/ml Nourseothricin Sulfate (clonNAT).

### Construction of gene knockout cassettes

Gene knockout cassettes consisted of three fragments, i.e., 2-kb homologous sequences of the 5′ and 3′ untranslated region (UTR) of the target gene flanking a nourseothricin acetyltransferase gene (*NAT1*). Two homologous sequences were designed to overlap with the *NAT1* gene. The three fragments were amplified separately, and these were then conjoined into complete cassettes by amplifying using nested primers targeting to both ends of the cassette.

### Confirmation of gene knockouts in mutants

To confirm knockout by PCR, we used two pairs of primers, each pair having one primer in the DNA flanking the targeted region to be knocked out and one in the *NAT1* gene, and amplified both intervening junctions. Another PCR reaction, using the two primers within the targeted region, was also performed to confirm the absence of the knockout gene, thereby ruling out the possibility of a duplication event.

Moreover, Southern blots were conducted on transformants to confirm knockout and to check if there had been any ectopic integrations of the drug-resistance cassette elsewhere in the genome.

### Rescue assay on EYR41_001410 in the TWF1042 mutant

Constructs were amplified from 2-kb upstream to 1-kb downstream of the EYR41_001410 sequence and fused with a G418 (Geneticin) drug cassette. We transformed 5 μg of constructs into 2.5×10^5^ of TWF1042 protoplasts and cultured them with 200 μg/ml of G418. Any resulting colonies were inoculated onto PDA plates with 150 μg/ml G418 for confirmation.

## Results

### Forward genetic screening identifies *A. oligospora* mutants with trapping defects

To establish a protocol for forward genetic screens that could identify genes involved in the trapping process of NTF, we optimized the mutagenesis conditions for a lethal dose (LD_50_) using EMS- or UV-treated protoplasts (6 seconds of 15 W UV or 12 μg/ml EMS). Then, the resulting 5552 mutagenized clones (1560 from UV- and 3992 from EMS-based mutagenesis) were isolated onto 48-well plates. We screened these 5552 mutant lines twice and identified 15 that exhibited defects in capturing *C. elegans* (Fig. 1). Unlike control wild-type strain TWF154, which formed numerous traps and captured all 30 *C. elegans* within 12 hours of our nematode-trapping assay, many live nematodes were still crawling over the mutant strains within the same timeframe and the mutants presented few or no traps (Fig. 2). Interestingly, some of the mutants exhibited delayed trap formation, with traps eventually being formed 24 hours after exposure to *C. elegans* (Supplementary figure 1). In summary, all of the mutants isolated from our genetic screens exhibit defects in trap development in the presence of nematode prey.

**Fig. 1.**
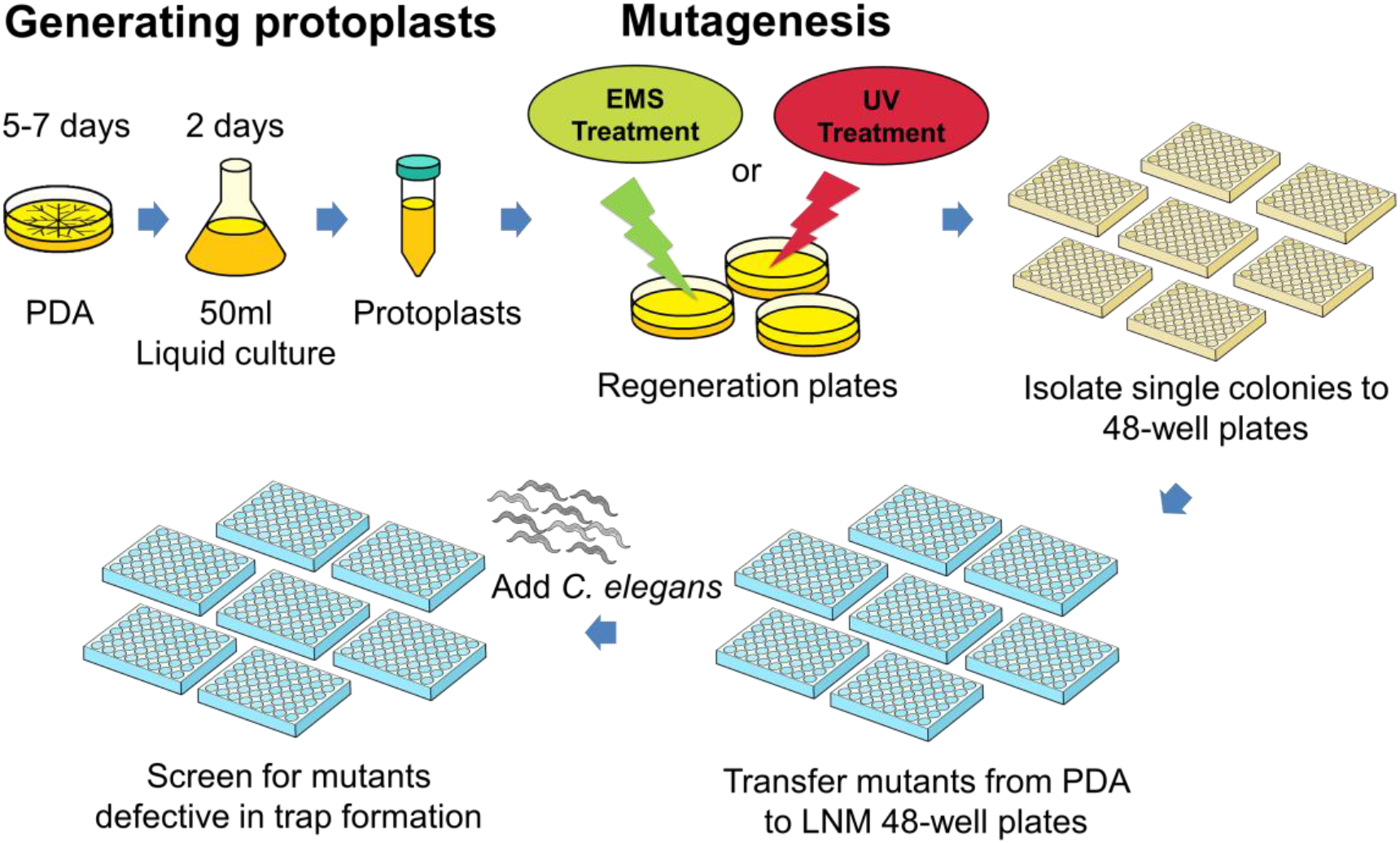
Schematic of our workflow for random mutagenesis and forward genetic screening of *A. oligospora*. Protoplasts were acquired from PDB liquid culture and were subsequently treated with EMS or UV for mutagenesis. The resulting colonies were separated out into 48-well PDA plates and screened on LNM plates. Mutants with trapping defects were selected after a rescreening process.

**Fig. 2.**
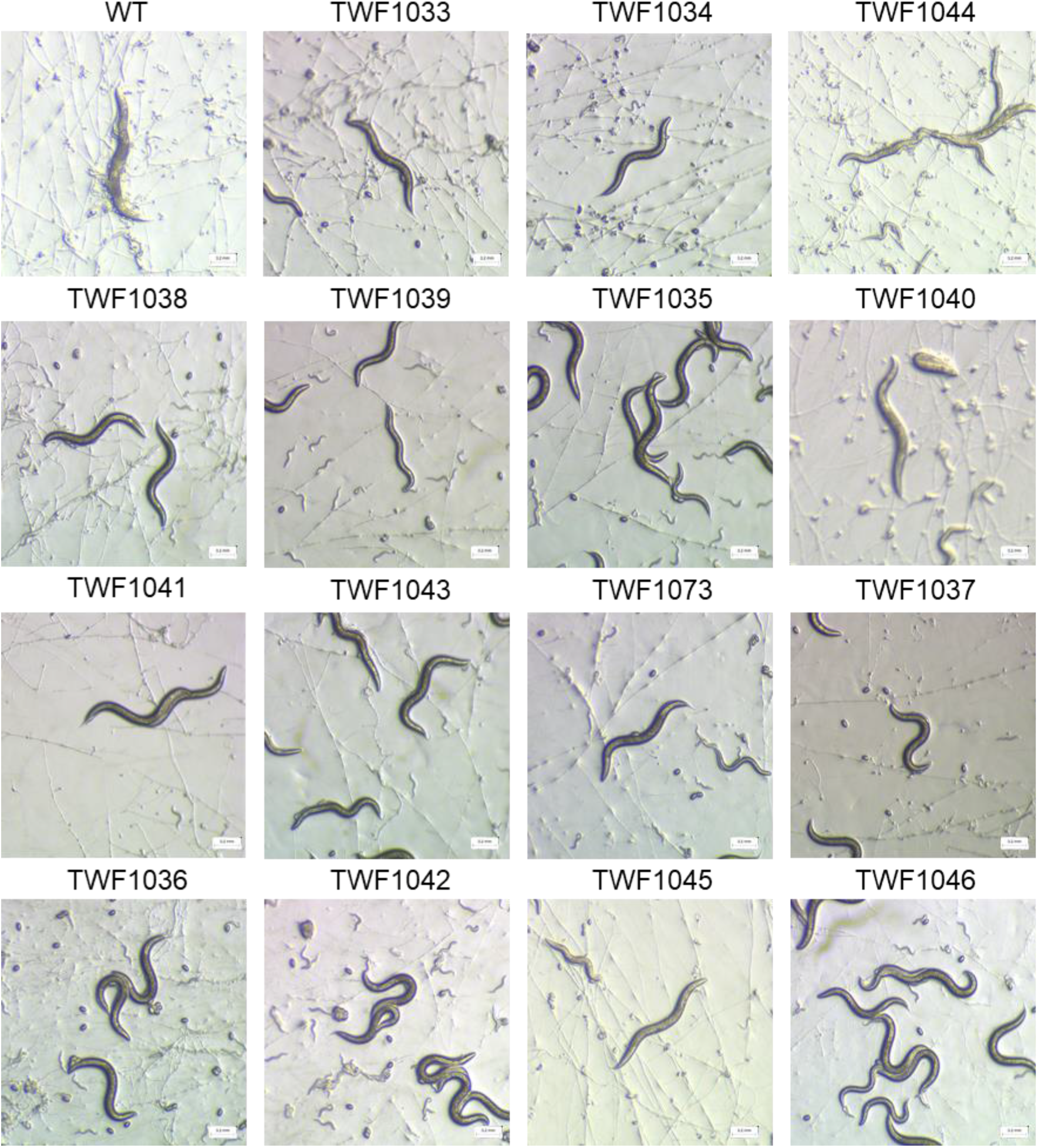
Random mutagenesis and genetic screening identified 15 *A. oligospora* mutants with defects in trap morphogenesis. All images are of *A. oligospora* upon 12-h exposure to N2 *C. elegans*.

### Phenotypic characterization of the 15 mutants exhibiting trapping defects

We further quantified their trap-forming capabilities in all 15 mutants that we had isolated from the genetic screens. We found that all mutant lines developed fewer traps relative to wild-type upon exposure to the same number of nematode prey, with trap morphogenesis being completely abolished in six of the mutant lines (Fig. 3A).

**Fig. 3.**
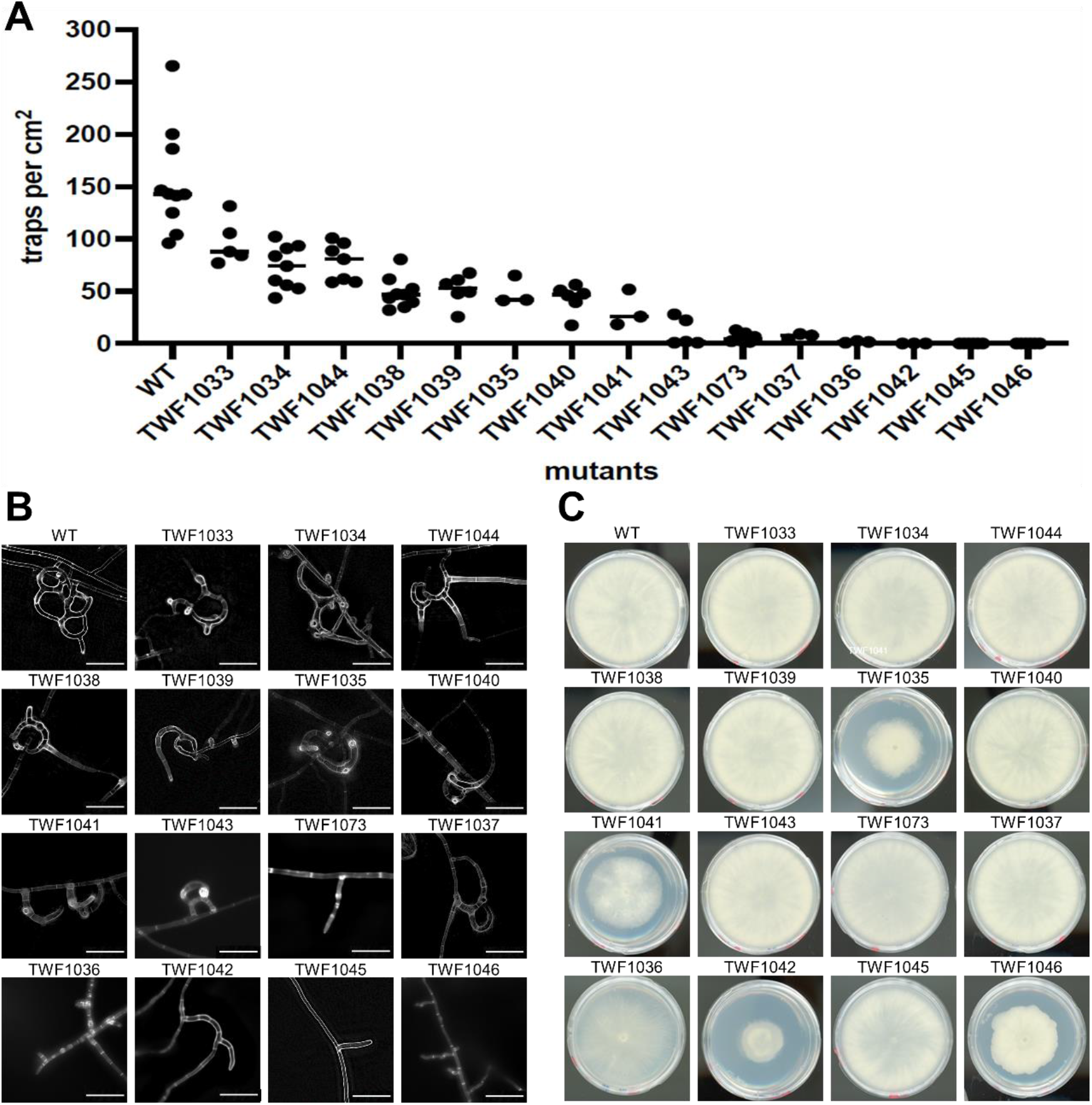
Phenotypic characterization of the 15 mutants identified from our forward genetic screen. (A) Quantification of trap numbers induced by *C. elegans* for the WT and 15 mutant lines. (B) Microscopic analyses of trap morphology for WT and the 15 mutant lines. Bars, 50 μm. (C) Growth of WT and 15 mutant lines on PDA plates (5-cm diameter) by day 5.

We then examined the morphology of the traps developed by these 15 mutants. The traps generated by wild-type *A. oligospora* are three-dimensional structures consisting of several loops of hyphae of varying size. In contrast, the traps formed by the mutants tended to have abnormal morphologies, either because the loops failed to fuse or the loops were irregularly shaped (Fig. 3B). It has been demonstrated previously that intercellular communication is critical for trap formation (Nordbring-Hertz et al., 2011; Youssar et al., 2019), so these phenotypes could result from deficient communication between hyphal cells or defects in initiating trap morphogenesis. Many of the mutants exhibited vegetative hyphal protrusions in response to nematode presence but were incapable of completing trap morphogenesis, some developed rudimentary traps, and others completely lacked any trap-like structures (Fig. 3B).

Next, we tested if these mutants exhibit general growth defects by culturing them on the rich medium potato dextrose agar (PDA). Four out of fifteen mutant lines showed growth defects relative to the wild-type strain, suggesting that these particular mutant lines harbor mutations in genes that may play pleotropic roles in general growth (Fig. 3C). The remaining 11 mutants displayed no overt growth differences compared to wild-type, implying that the mutated genes might play more specific roles in *A. oligospora* trap formation.

### Whole genome sequencing analysis of the 15 mutant lines identifies potential candidate genes involved in trap morphogenesis

To identify the mutations in the genomes of the 15 mutant lines, we conducted whole genome sequencing and remapped all sequencing data to the wild-type TWF154 reference genome. On average, 2700 mutations encompassing noncoding and coding sequences were identified in each of these mutants, but many of those mutations were common to all mutant lines, perhaps representing background mutations between the mutagenized clone and the original sequenced clone (Supplementary figure 2). Such mutations were ruled out from further analyses (Table 1). Of the 10-89 mutations remaining in each of the mutant lines, we focused on those occurring within exonic sequences to further refine the group of candidate genes (Table 1). In certain mutant lines, such as TWF1037, TWF1042, TWF1046 and TWF1073, we identified mutations in exons of more than 20 genes, whereas for two mutant strains (TWF1033 and TWF1034) we did not identify any coding genes with mutations, indicating that the mutations that caused trapping defects in these two latter mutants likely occur in noncoding regions of the genome.

**Table 1.** Summary of numbers of mutations (after filtering out common mutations) among the 15 mutant lines, as identified by whole genome sequencing. Exonic mutations encompass frameshift indels, non-frameshift indels, non-synonymous, synonymous and stop-gain mutations. Indel, insertion/deletion; SNV, single nucleotide variation.

| Mutant | Upstream | Downstream | Intergenic | Intronic | 5'UTR | 3'UTR | Exonic | Frameshift<br>indels | Non-frameshift<br>indelss | Non-<br>synonymous<br>SNV | Synonymous<br>SNV | Stop-gain | Total |
| --- | --- | --- | --- | --- | --- | --- | --- | --- | --- | --- | --- | --- | --- |
| TWF1033 | 1 | 3 | 6 | 1 | 0 | 0 | 0 | 0 | 0 | 0 | 0 | 0 | 11 |
| TWF1034 | 1 | 2 | 5 | 2 | 0 | 0 | 0 | 0 | 0 | 0 | 0 | 0 | 10 |
| TWF1035 | 4 | 4 | 7 | 4 | 1 | 0 | 3 | 1 | 0 | 2 | 0 | 0 | 23 |
| TWF1036 | 5 | 8 | 9 | 2 | 2 | 4 | 8 | 1 | 0 | 6 | 1 | 0 | 38 |
| TWF1037 | 10 | 6 | 28 | 5 | 9 | 7 | 24 | 0 | 0 | 18 | 6 | 0 | 89 |
| TWF1038 | 5 | 6 | 8 | 7 | 2 | 2 | 12 | 0 | 0 | 6 | 5 | 1 | 42 |
| TWF1039 | 2 | 5 | 10 | 0 | 2 | 1 | 10 | 0 | 0 | 7 | 3 | 0 | 30 |
| TWF1040 | 5 | 7 | 5 | 1 | 2 | 1 | 10 | 0 | 0 | 7 | 3 | 0 | 31 |
| TWF1041 | 3 | 5 | 9 | 4 | 4 | 1 | 17 | 1 | 0 | 12 | 4 | 1 | 43 |
| TWF1042 | 13 | 3 | 15 | 8 | 4 | 0 | 26 | 3 | 0 | 14 | 9 | 0 | 69 |
| TWF1043 | 6 | 7 | 11 | 10 | 2 | 2 | 12 | 0 | 0 | 6 | 5 | 1 | 50 |
| TWF1044 | 5 | 2 | 6 | 7 | 4 | 2 | 3 | 0 | 0 | 2 | 1 | 0 | 29 |
| TWF1046 | 14 | 3 | 12 | 11 | 9 | 5 | 26 | 2 | 0 | 9 | 12 | 3 | 80 |
| TWF1073 | 28 | 6 | 4 | 6 | 14 | 4 | 22 | 9 | 3 | 4 | 1 | 5 | 84 |

In the final step of our mutation selection process, we excluded genes having synonymous mutations, and selected genes that gained a misplaced stop codon for further functional studies (Table 1). We identified loss-of-function mutations, such as stop gain mutations, frameshift indels, and nonsynonymous SNV, among the mutants (Fig. 4). We used gene ontology analysis to predict the functions of the mutated genes identified from our sequencing analyses and discovered that they play roles in signaling, transcription or membrane transport (Supplementary Table 2). Together, these analyses have revealed a set of genes that, when mutated, may contribute to the phenotype of impaired trap formation and nematode predation.

**Fig. 4.**
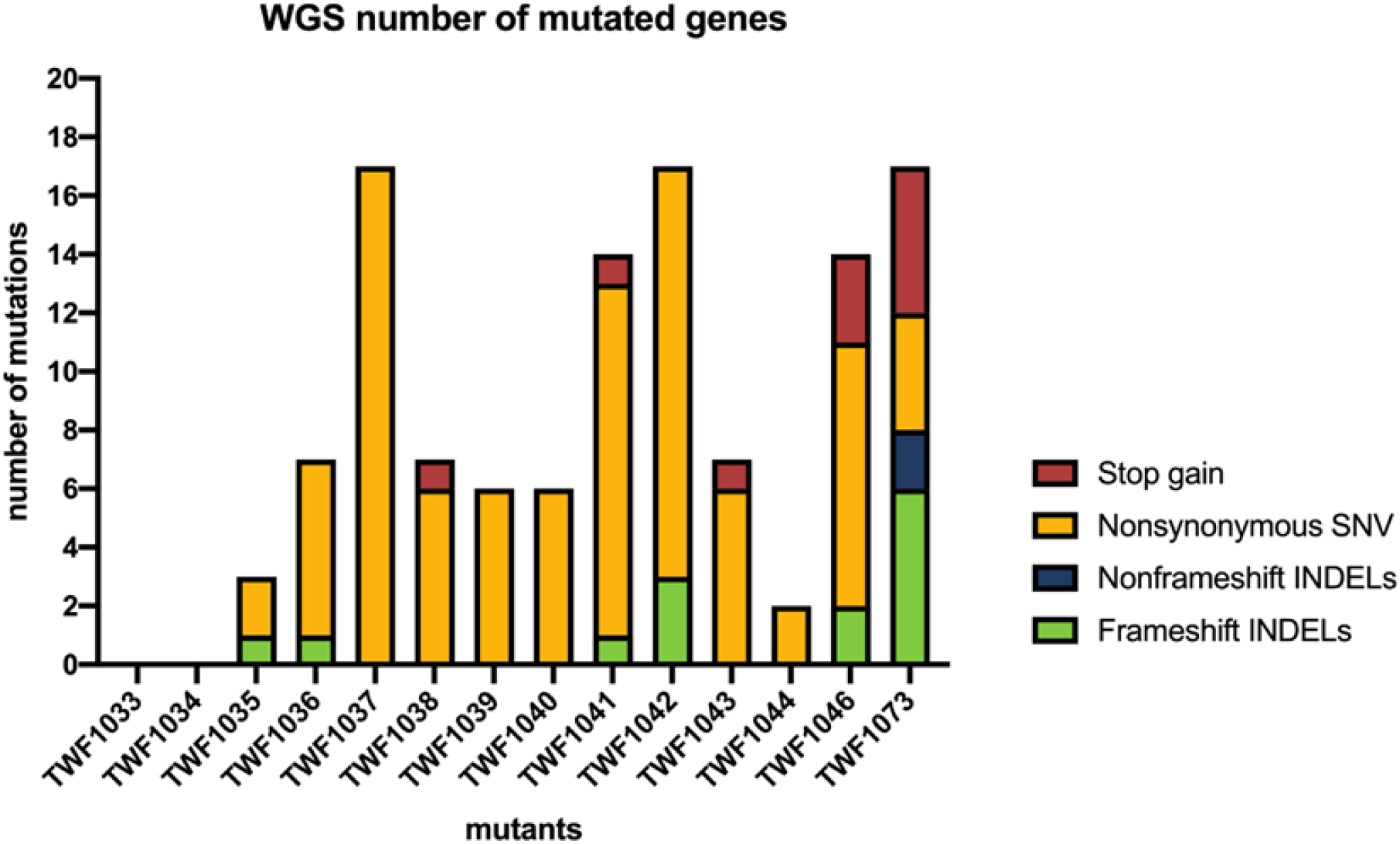
Numbers of mutated genes (after filtration procedures) in each mutant. Different colors represent different types of exonic mutations (excluding synonymous mutations).

### Frameshift indel mutation in gene EYR41_001410 induces phenotypic defects in the TWF1042 mutant strain

To further assess identified candidate mutated genes and establish which mutations were causative for the phenotypes observed in the mutants, we focused on mutant strain TWF1042 that had mutations in seventeen candidate genes, including three frameshift indel mutations (Fig. 4). Reasoning that these three frameshift mutations likely abolished the respective protein function, we examined our gene ontology predictions and discovered that one putative sequence had a protein kinase domain (EYR41_005093), another had a YAP-binding/ALF4/Glomulin domain (EYR41_001410), and the other was of unknown function (EYR41_008629). In *Saccharomyces cerevisiae*, YAP-binding proteins function in responses to oxidative stress (Gulshan et al., 2004). In *Arabidopsis thaliana*, Aberrant root formation protein 4 (Alf4) is involved in the initiation of lateral root formation (DiDonato et al., 2004). Deletion of the *Glomulin* gene was reported to affect differentiation in vascular smooth muscle cells in mouse (Arai et al., 2003). Given these diverse biological functions of the YAP-binding/ALF4/Glomulin protein family, we hypothesized that the protein encoded by *EYR41_001410* in *A. oligospora*, which we have named *YBP1* (<u>Y</u>AP-<u>b</u>inding protein 1), might play an important role in hyphal growth in this fungus. *YPB1* appears to be a rapidly-evolving gene. It shares ~70% protein sequence identity among NTF, but only ~20% protein sequence identity with other ascomycetes (Fig. 5A). Consequently, the YAP-binding protein gene family may play diverse roles in a variety of fungal spices.

**Fig. 5.**
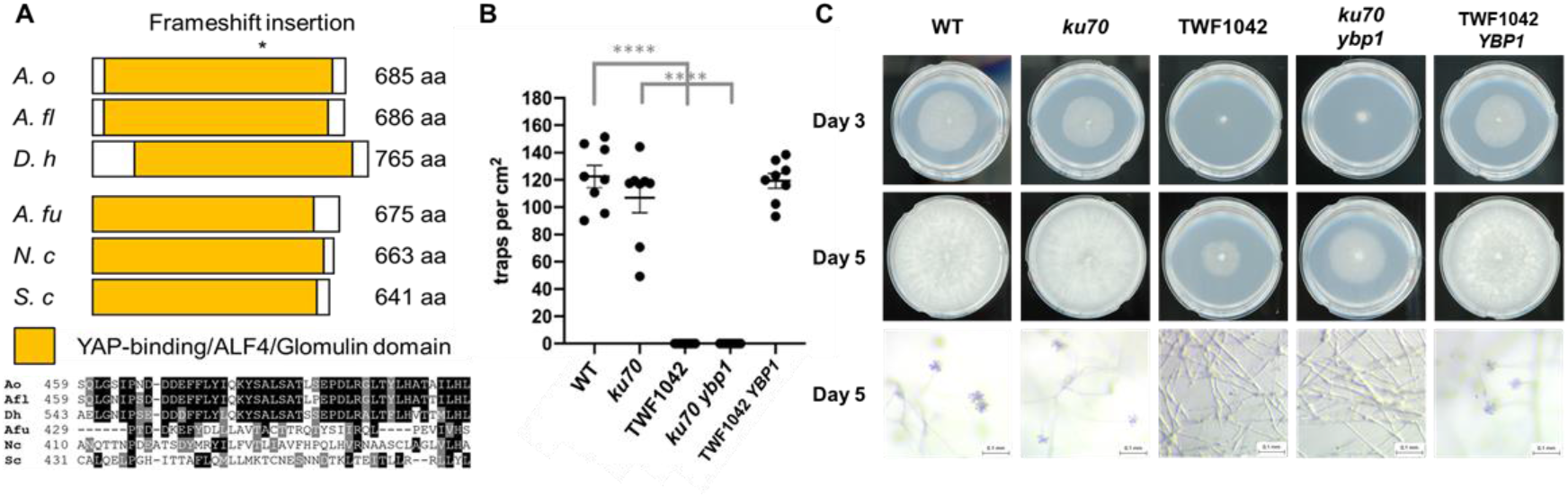
Mutations in *YBP1* cause the phenotypic defects in growth, trap morphogenesis and conidiation observed in the randomly-mutagenized strain TWF1042. (A) Schematic representation of the domain structure and partial sequence alignment of *A. oligospora* YBP1 and related fungal homologs. *A. oligospora* (*A. o*), *Arthrobotrys flagrans* (*A. fl*), *Dactylellina haptotyla* (*D. h*), *Aspergillus fumigatus* (*A. fu*), *Neurospora crassa* (*N. c*), *Saccharomyces cerevisiae* (*S. c*). Asterisk represents the site where frameshift insertion was found. (B) Quantification of trap numbers induced by *C. elegans* presence for the WT, *ku70*, TWF1042, *ku70 ybp1*, and a TWF1042-*YBP1* rescue strain. (C) Representative images of growth (day 3), aerial hyphae (day 5), and conidiation (day 5) for the WT, *ku70*, TWF1042, *ku70 ybp1*, *and* TWF1042*-YBP1* rescue strain. Colonies were grown on PDA plates (5-cm diameter).

To study the function of *YPB1*, we constructed a gene deletion mutant via homologous recombination and examined the phenotypes of the resulting *ypb1* mutant. We found that *ypb1* displayed phenotypes similar to those exhibited by our randomly mutagenized strain, TWF1042, including slow growth, lack of conidiation and severe defects in trap formation (Fig. 5B and C). These results indicate that the frameshift insertion mutation in *YPB1* likely caused the phenotypic defects observed in strain TWF1042. To validate that supposition, we expressed the wild-type allele of *YPB1* under its endogenous promoter in TWF1042 to examine if addition of a wild-type copy of *YPB1* could rescue the phenotypic defects displayed by TWF1042. Indeed, addition of the wild-type *YPB1* allele to TWF1042 complemented its defects in trap formation, growth, and conidiation, demonstrating that the frameshift insertion mutation of *YPB1* caused the phenotypic defects observed for TWF1042 (Fig. 5C).

## Discussion

In this study, we established a protocol to conduct forward genetic screens in the NTF, *A. oligospora*. We used EMS or UV to mutagenize protoplasts and screened out fifteen mutants exhibiting defects in trap formation. Subsequent whole genome sequencing identified candidate genes harboring mutations in these mutants. Finally, we demonstrate that a frameshift mutation of the YAP-binding/ALF4/Glomulin domain-containing gene, *YPB1*, caused the nematode-trapping defects observed in one of the randomly-mutagenized mutant strains, TWF1042.

Although we identified 15 mutants with defects in trap morphogenesis from our genetic screens, we consider the success rate in isolating mutants to be low (15 out of ~5,500 mutagenized clones), and our screens were far from achieving mutagenesis saturation. We believe that two factors contributed to this result. First, mutagen dosages may have been too light and, second, some of our mutagenized clones could be heterokaryons of mixed genetic backgrounds, which could mask phenotypes caused by mutations. We purposely used lower mutagen dosages because laboratory methods for conducting genetic crosses of *A. oligospora* have yet to be established. If too many mutations had been generated in the background genome, it would be challenging to identify the mutations causing the observed phenotypes without undertaking genetic mapping analyses. Since the hyphae of *A. oligospora* contain multiple nuclei, it is possible that a small proportion of the protoplasts we generated and mutagenized harbored more than one nucleus, and it is also possible that some of the single mycelium colonies we isolated had fused with a neighboring colony of a different genetic background. Both scenarios could lead to heterokaryons occurring in our *A. oligospora* mutant libraries.

We believe that to make forward genetic screening even more applicable to NTF study, laboratory genetic crosses must be established. Doing so would enable significantly higher mutagen dosages to be applied, rendering mutant identification more efficient. In summary, we have established a method for conducting random mutagenesis in a non-model fungus, followed by resequencing of the mutants to identify candidate genes contributing to observed phenotypes. We have revealed that YBP1 plays a critical role in the physiology and development of *A. oligospora* and also identified several other candidate genes in which mutations might cause defects in trap morphogenesis. We envisage that our methodology could facilitate future genetic studies in other enigmatic fungi.

## Acknowledgments

The authors thank the IMB genomic core for conducting Illumina sequencing and John O’Brien’s comments on the manuscript. This work was supported by Academia Sinica Career Development Award AS-CDA-106-L03 and Taiwan Ministry of Science and Technology grant 106-2311-B-001-039-MY3 to YPH.

---

**Supplementary fig 1.**
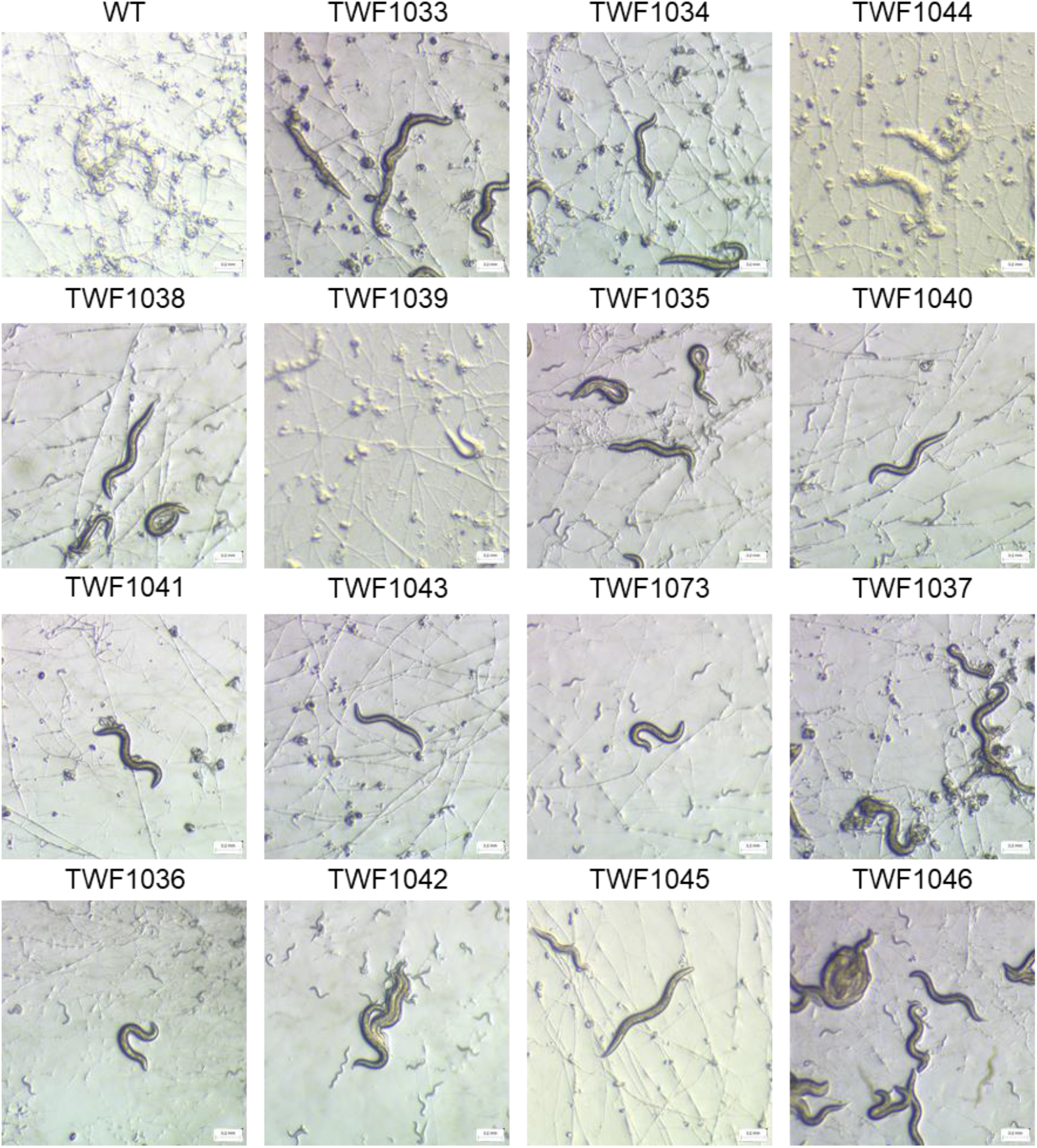
Random mutagenesis and forward genetic screening identified 15 mutants with defects in capturing *C. elegans*. *C. elegans* were paralyzed and dead in WT group but there were still living worms in mutants. Micrographs were captured at the 24-h time-point. Some mutants showed delayed trap formation and others still failed to form traps.

**Supplementary fig 2.**
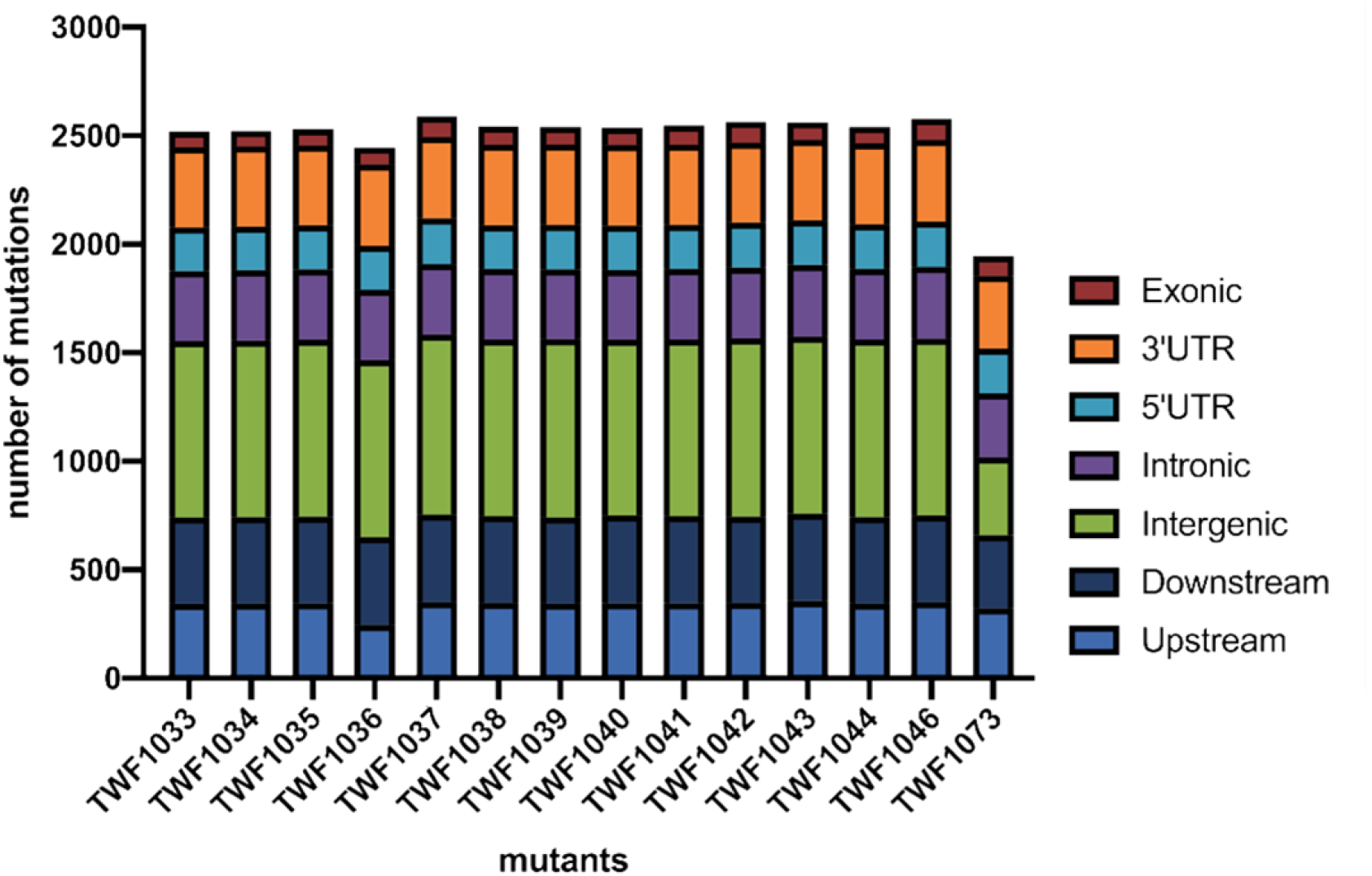
Total mutations annotated for each mutant. Different types of mutation are individually colored.

**Supplementary Table 2.** Information on selected genes (after filtration procedures) to conduct targeted gene knockout. “Chr” column presents the chromosome that the mutations were located. “Start” and “End” column present the exact sites where the mutations were found. “Ref” and “Alt” column present the nucleotides from reference genome and mutants respectively. “Exonic Func.” presents the type of the mutations. “GO terms” presents the predicted function of the gene.

| Mutant | Gene | Chr | Start | End | Ref | Alt | Exonic Func. | GO terms |
| --- | --- | --- | --- | --- | --- | --- | --- | --- |
| TWF1035 | EYR41_000049 | scaffold_1 | 149046 | 149046 | - | TCTGTAA | frameshift insertion | Creatinase/aminopeptidase-like |
| TWF1036 | EYR41_003391 | scaffold_2 | 2818724 | 2818724 | A | G | nonsynonymous SNV | Major facilitator, sugar transporter-like |
| TWF1036 | EYR41_009094 | scaffold_6 | 793179 | 793179 | T | A | nonsynonymous SNV | Zinc finger, CCCH-type superfamily |
| TWF1037 | EYR41_002645 | scaffold_2 | 399109 | 399109 | T | A | nonsynonymous SNV | F-box-like domain superfamily |
| TWF1037 | EYR41_008023 | scaffold_5 | 1235392 | 1235392 | T | A | nonsynonymous SNV | Protein kinase domain |
| TWF1038 | EYR41_005196 | scaffold_3 | 1505468 | 1505468 | - | TA | stop-gain | MFS transporter superfamily |
| TWF1038 | EYR41_011609 | scaffold_8 | 2469219 | 2469219 | T | A | nonsynonymous SNV | Cytochrome P450, E-class, group I |
| TWF1039 | EYR41_008594 | scaffold_5 | 3072546 | 3072546 | G | A | nonsynonymous SNV | Zinc finger, GATA-type;zinc finger, GATA-type |
| TWF1041 | EYR41_005971 | scaffold_3 | 3955532 | 3955532 | A | T | stop-gain | Copper amine oxidase, catalytic domain |
| TWF1042 | EYR41_001410 | scaffold_1 | 4520093 | 4520093 | - | TTCCGACGAATTT | frameshift insertion | YAP-binding/ALF4/Glomulin |
| TWF1042 | EYR41_005093 | scaffold_3 | 1186858 | 1186858 | - | T | frameshift insertion | Protein kinase domain |
| TWF1042 | EYR41_008629 | scaffold_5 | 3181771 | 3181771 | T | - | frameshift deletion | Protein of unknown function DUF4246 |
| TWF1043 | EYR41_005196 | scaffold_3 | 1505468 | 1505468 | - | TA | stop-gain | MFS transporter superfamily |
| TWF1046 | EYR41_003064 | scaffold_2 | 1780257 | 1780257 | A | C | stop-gain | Velvet domain |
| TWF1046 | EYR41_010493 | scaffold_7 | 1849994 | 1849994 | A | T | nonsynonymous SNV | Domain of unknown function DUF2183 |

## References

1. Arai, T., Kasper, J.S., Skaar, J.R., Ali, S.H., Takahashi, C., and DeCaprio, J.A.J.P.o.t.N.A.o.S. (2003). Targeted disruption of p185/Cul7 gene results in abnormal vascular morphogenesis. Proceedings of the National Academy of Sciences 100, 9855–9860.

2. Bolger, A.M., Lohse, M., and Usadel, B.J.B. (2014). Trimmomatic: a flexible trimmer for Illumina sequence data. Bioinformatics 30, 2114–2120.

3. DiDonato, R.J., Arbuckle, E., Buker, S., Sheets, J., Tobar, J., Totong, R., Grisafi, P., Fink, G.R., and Celenza, J.L.J.T.P.J. (2004). Arabidopsis ALF4 encodes a nuclear‐localized protein required for lateral root formation. The Plant Journal 37, 340–353.

4. Ellegren, H.J.T.i.e., and evolution (2014). Genome sequencing and population genomics in non-model organisms. Trends in ecology & evolution 29, 51–63.

5. Gray, N.F. (1983). Ecology of nematophagous fungi: Distribution and habitat. Annals of Applied Biology 102, 501–509.

6. Gulshan, K., Rovinsky, S.A., and Moye-Rowley, W.S.J.E.c. (2004). YBP1 and its homologue YBP2/YBH1 influence oxidative-stress tolerance by nonidentical mechanisms in Saccharomyces cerevisiae. Eukaryot Cell 3, 318–330.

7. Hsueh, Y.P., Gronquist, M.R., Schwarz, E.M., Nath, R.D., Lee, C.H., Gharib, S., Schroeder, F.C., and Sternberg, P.W. (2017). Nematophagous fungus *Arthrobotrys oligospora* mimics olfactory cues of sex and food to lure its nematode prey. eLife 6.

8. Hsueh, Y.P., Mahanti, P., Schroeder, F.C., and Sternberg, P.W. *Current Biology* (2013). Nematode-trapping fungi eavesdrop on nematode pheromones. Curr Biol 23, 83–86.

9. Li, H., and Durbin, R.J.b. (2009). Fast and accurate short read alignment with Burrows–Wheeler transform. Bioinformatics 25, 1754–1760.

10. Li, H.J.a.p.a. (2013). Aligning sequence reads, clone sequences and assembly contigs with BWA-MEM. arXiv:1303.3997 [q-bio.GN]

11. Li, H.J.B. (2011). A statistical framework for SNP calling, mutation discovery, association mapping and population genetical parameter estimation from sequencing data. Bioinformatics 27, 2987–2993.

12. Li, J., Wu, R., Wang, M., Borneman, J., Yang, J., and Zhang, K.-Q.J.F.b. (2019). The pH sensing receptor AopalH plays important roles in the nematophagous fungus *Arthrobotrys oligospora*. Fungal biology 123, 547–554.

13. Li, X., Kang, Y.-Q., Luo, Y.-L., Zhang, K.-Q., Zou, C.-G., and Liang, L.-M.J.J.o.M. (2017). The NADPH oxidase AoNoxA in *Arthrobotrys oligospora* functions as an initial factor in the infection of *Caenorhabditis elegans*. Journal of Microbiology 55, 885–891.

14. Lopez-Llorca, L.V., Maciá-Vicente, J.G., and Jansson, H.B. (2007). Mode of Action and Interactions of Nematophagous Fungi. *In Integrated management and biocontrol of vegetable and grain crops nematodes*. Springer, Dordrecht 2, 51–76.

15. Nordbring-Hertz, B., Jansson, H.-B., and Tunlid, A. (2011). Nematophagous fungi.

16. Poplin, R., Ruano-Rubio, V., DePristo, M.A., Fennell, T.J., Carneiro, M.O., Van der Auwera, G.A., Kling, D.E., Gauthier, L.D., Levy-Moonshine, A., and Roazen, D.J.B. (2017). Scaling accurate genetic variant discovery to tens of thousands of samples. bioRxiv, 201178.

17. Russell, J.J., Theriot, J.A., Sood, P., Marshall, W.F., Landweber, L.F., Fritz-Laylin, L., Polka, J.K., Oliferenko, S., Gerbich, T., and Gladfelter, A.J.B.b. (2017). Non-model model organisms. BMC biology 15, 1–31.

18. Wang, K., Li, M., and Hakonarson, H.J.N.a.r. (2010). ANNOVAR: functional annotation of genetic variants from high-throughput sequencing data. Nucleic Acids Research 38, e164–e164.

19. Wang, R., Wang, J., and Yang, X. (2015). The extracellular bioactive substances of *Arthrobotrys oligospora* during the nematode-trapping process. Biological Control 86, 60–65.

20. Wu, B., Hussain, M., Zhang, W., Stadler, M., Liu, X., and Xiang, M.J.M. (2019). Current insights into fungal species diversity and perspective on naming the environmental DNA sequences of fungi. Mycology 10, 127–140.

21. Yang, C.T., Vidal-Diez de Ulzurrun, G., Goncalves, A.P., Lin, H.C., Chang, C.W., Huang, T.Y., Chen, S.A., Lai, C.K., Tsai, I.J., Schroeder, F.C., et al. (2020). Natural diversity in the predatory behavior facilitates the establishment of a robust model strain for nematode-trapping fungi. Proceedings of the National Academy of Sciences U S A.

22. Youssar, L., Wernet, V., Hensel, N., Yu, X., Hildebrand, H.-G., Schreckenberger, B., Kriegler, M., Hetzer, B., Frankino, P., and Dillin, A.J.P.g. (2019). Intercellular communication is required for trap formation in the nematode-trapping fungus *Duddingtonia flagrans*. Plos Genetics 15, e1008029.

23. Zhen, Z., Xing, X., Xie, M., Yang, L., Yang, X., Zheng, Y., Chen, Y., Ma, N., Li, Q., Zhang, K.-Q.J.F.G., et al. (2018). MAP kinase Slt2 orthologs play similar roles in conidiation, trap formation, and pathogenicity in two nematode-trapping fungi. Fungal Genetics and Biology 116, 42–50.

